# A bipartite chromatophore targeting peptide and N-terminal processing of proteins in the *Paulinella* chromatophore

**DOI:** 10.1101/2021.09.19.461000

**Authors:** Linda Oberleitner, Andreas Perrar, Luis Macorano, Pitter F. Huesgen, Eva C. M. Nowack

## Abstract

The cercozoan amoeba *Paulinella chromatophora* contains photosynthetic organelles - termed chromatophores - that evolved from a cyanobacterium ∼100 million years ago, independently from plastids in plants and algae. Despite its more recent origin, at least one third of the chromatophore proteome consists of nucleus-encoded proteins that are imported by an unknown mechanism across the chromatophore double envelope membranes. Chromatophore-targeted proteins fall into two classes. Proteins exceeding 250 amino acids carry a conserved N-terminal sequence extension, termed the ‘chromatophore transit peptide’ (crTP), that is presumably involved in guiding these proteins into the chromatophore. Short imported proteins do not carry discernable targeting signals. To explore whether the import of protein is accompanied by their N-terminal processing, here we used a mass spectrometry-based approach to determine protein N-termini in *Paulinella chromatophora* and identified N-termini of 208 chromatophore-localized proteins. Our study revealed extensive N-terminal modifications by acetylation and proteolytic processing in both, the nucleus and chromatophore-encoded fraction of the chromatophore proteome. Mature N-termini of 37 crTP-carrying proteins were identified, of which 30 were cleaved in a common processing region. Our results imply that the crTP mediates trafficking through the Golgi, is bipartite and surprisingly only the N-terminal third (‘part 1’) becomes cleaved upon import, whereas the rest (‘part 2’) remains at the mature proteins. In contrast, short imported proteins remain largely unprocessed. Finally, this work sheds light on N-terminal processing of proteins encoded in an evolutionary-early-stage photosynthetic organelle and suggests host-derived post-translationally acting factors involved in dynamic regulation of the chromatophore-encoded chromatophore proteome.

**One sentence summary:** Proteins targeted to the evolutionary-early-stage photosynthetic organelle of *Paulinella* carry a bipartite N-terminal targeting sequence that is only partially removed upon protein import.

## Introduction

Besides mitochondria and primary plastids that evolved via endosymbioses more than one billion years ago, recently, a third organelle of primary endosymbiotic origin has been identified (Nowack, 2014; Gabr et al., 2020). The photosynthetic ‘chromatophore’ of cercozoan amoebae of the genus *Paulinella* evolved around 100 million years ago from a cyanobacterium (Marin et al., 2005; Delaye et al., 2016). Following establishment of the endosymbiosis the chromatophore genome was reduced to around one third of its original size and many lost functions are compensated by the import of nucleus-encoded proteins (Nowack and Grossman, 2012; Singer et al., 2017; Oberleitner et al., 2020). In previous studies, we identified together >500 nucleus-encoded, chromatophore-targeted proteins in *P. chromatophora* (Singer et al., 2017; Oberleitner et al., 2020). These proteins comprise more than one third of the chromatophore proteome and fall into two classes: short imported proteins [<90 amino acids (aa)] that lack obvious targeting signals, and long imported proteins (>250 aa) that carry a conserved N-terminal sequence extension (Singer et al., 2017), referred to as the ‘chromatophore transit peptide’ (crTP).

In plants and algae, nucleus-encoded proteins that are targeted to primary plastids are synthesized on eukaryotic ribosomes as pre-proteins carrying an N-terminal chloroplast transit peptide (cTP). The unfolded pre-proteins are bound by specific chaperones in the cytosol and guided by the cTP to the TIC/TOC translocon. Upon import, the cTP is cleaved by the Stromal Processing Peptidase (SPP) (Teixeira and Glaser, 2013). Mature stromal proteins can now fold into their functional conformation. Nucleus-encoded proteins designated to the thylakoid lumen reach their final destination via the bacterial Sec- or Tat-pathways. This usually requires a bipartite targeting signal in which the cTP is followed by a bacterial signal peptide (Schünemann, 2007). In addition to the import-related processing of nucleus-encoded proteins, many chloroplast-encoded proteins can be N-terminally modified, which regulates targeting, stability and function (Giglione and Meinnel, 2001; Huesgen et al., 2013; Rowland et al., 2015; Varland et al., 2015; Linster and Wirtz, 2018). These modifications include N-terminal cleavage or trimming by proteases, excision of the initiating methionine (iMet) by methionine aminopeptidases, or N-acetylation by N-acetyltransferases.

In *P. chromatophora*, the mechanisms that underlie import of long and short nucleus-encoded proteins into the chromatophore are largely unknown. Whether import involves N-terminal processing of the imported proteins has not been studied yet. A translocon similar to the TIC/TOC complex seems to be missing. The only orthologs of components of this multiprotein complex found in *P. chromatophora* are those of Tic21 [possibly a protein conducting channel in the inner envelope, but its function is debated (Duy et al., 2007)] and the regulatory components Tic32 and Tic62; orthologs of Tic110, Tic20, and Toc75 which form the major transport channels through inner and outer membrane were not identified (Gagat and Mackiewicz, 2014). It has been shown experimentally, that short imported proteins without a crTP are synthesized on eukaryotic ribosomes and one of them, the photosystem I subunit PsaE, was detected in the Golgi by immunogold electron microscopy, suggesting the Golgi as transport intermediate en route to the chromatophore (Nowack and Grossman, 2012). Whether the same applies for crTP-carrying long proteins is unknown. With approximately 200 aa, the crTP is unusually long for a targeting sequence. A signal peptide that would direct the protein to the secretory pathway is neither predicted at the N-terminus of short nor long imported proteins using SignalP (Almagro Armenteros et al., 2019). The crTP shows no similarity to other known proteins or protein domains that could provide a hint towards its function but a high degree of sequence conservation between chromatophore-targeted proteins including those in the photosynthetic sister species *P. micropora* (Singer et al., 2017; Lhee et al., 2021).

The facts that a method for the genetic manipulation of *P. chromatophora* has not been established yet and despite several attempts no antibodies specific against any long imported proteins could be raised, have severely hampered the dissection of the protein import process into the chromatophore. Here we used an unbiased mass spectrometry-based method, termed High-efficiency Undecanal-based N-Termini EnRichment (HUNTER) (Weng et al., 2019), to obtain experimental data on chromatophore protein N-termini and thereby elucidate N-terminal protein processing in the chromatophore, including processing events that are related to protein import. Our data reveals abundant N-terminal modifications of both nucleus and chromatophore-encoded chromatophore-localized proteins. Most importantly, it suggests that the crTP of long imported proteins is bipartite and surprisingly only partially removed from the mature protein upon import. Together with a bioinformatic characterization of common motifs and conserved structural elements within the crTP, our results suggest an import mechanism for crTP-carrying proteins that involves the fusion of Golgi-derived clathrin-coated vesicles with the outer chromatophore envelope membrane.

## Results

### Identification of Protein N-termini in the Chromatophore by HUNTER

We extracted proteins from isolated chromatophores and enriched N-terminal peptides using HUNTER (**Fig. 1A**). In short, protein N-termini were modified by reductive dimethylation before proteome digestion using trypsin. This is followed by reaction with the long-chain aldehyde undecanal, which modified all peptides carrying a free trypsin-generated primary amine. N-terminal peptides remain inert because they were either modified in the dimethylation reaction or endogenously blocked by acetylation, allowing their specific enrichment by C18 solid phase extraction-mediated depletion of the highly hydrophobic undecanal-modified internal and C-terminal peptides. Enriched N-terminal peptides were identified by high-resolution nano-flow liquid chromatography coupled to tandem mass spectrometry (nLC-MS/MS) using a database containing all protein models derived from *P. chromatophora* nuclear transcripts and the chromatophore genome.

**Figure 1.**
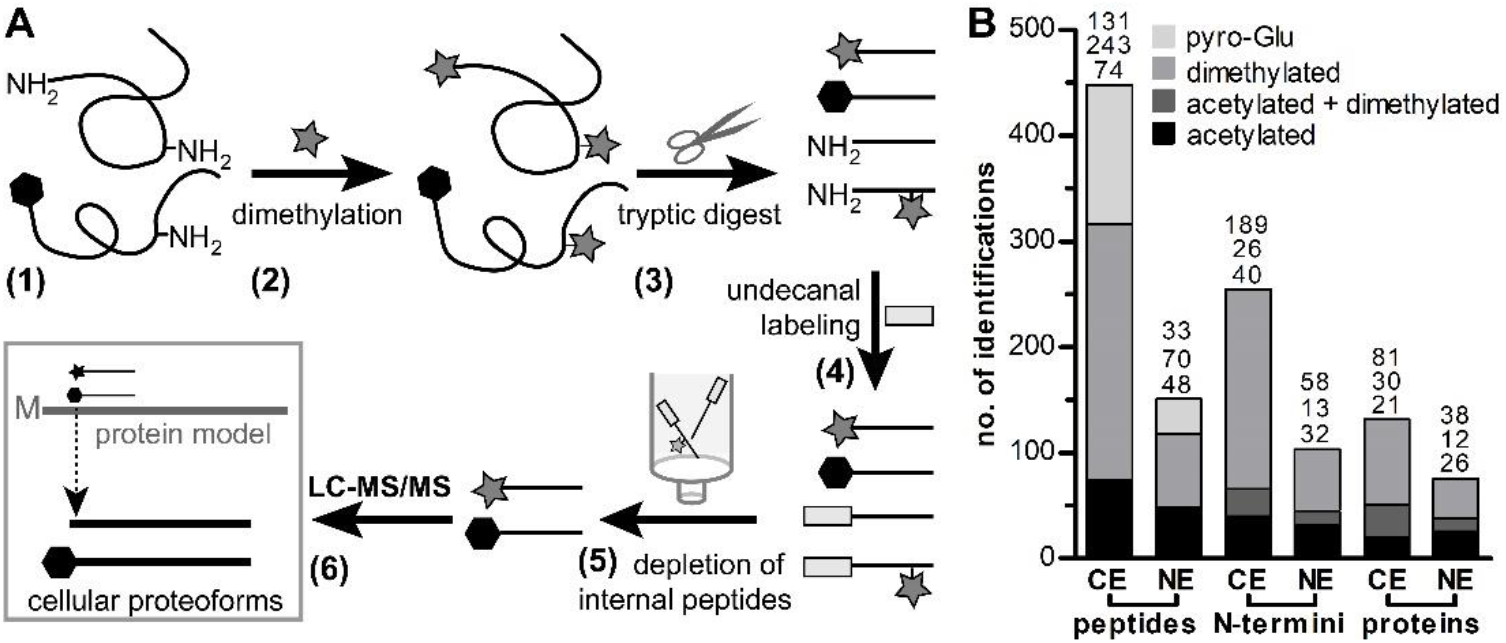
Application of HUNTER for the identification of N-termini in the chromatophore. **(A)** Schematic representation of the HUNTER workflow: (1) Proteins with naturally acetylated (hexagon) or free N-termini are purified from isolated chromatophores; (2) free N-terminal α- and lysine ε-amines are dimethylated (grey stars); (3) proteome is digested with trypsin; (4) free α-amines resulting from digestion are modified with undecanal (light grey rectangles); (5) undecanal-modified peptides are depleted on a reverse-phase column; (6) enriched N-terminal peptides are analyzed by nLC-MS/MS. **(B)** Total numbers of N-terminal peptides, corresponding unique N-termini, and corresponding proteins identified in triplicates of chromatophore lysates. A color code indicates the number of N-termini or proteins represented by acetylated, dimethylated, or both kinds of peptides sorted by chromatophore (CE) and nucleus-encoded (NE) proteins.

In total, this approach identified 599 peptides in chromatophore lysates at an FDR <0.01, of which 313 were dimethylated indicating N-termini with free primary amines *in vivo*, 122 from endogenously modified acetylated N-terminal peptides, and 164 pyro-glutamate modified peptides (**Fig. 1B**). As pyro-Glu modification may arise from endogenous modification, but also from spontaneous N-terminal glutamine cyclisation after tryptic digest, these peptides were not further considered (Demir et al., 2021). Of the remaining 435 (acetylated or dimethylated) bona fide N-terminal peptides, 317 were derived from chromatophore and 118 from nucleus-encoded proteins. We then further summarized N-terminal peptides that differed only by their N-terminal modification (i.e., acetylated and dimethylated versions of the same peptide) or only by their C-termini (i.e., resulting from missed trypsin cleavage sites), resulting in 255 and 103 unique N-termini, derived from 132 chromatophore and 76 nucleus-encoded proteins, respectively (**Fig. 1B; Table S1**). Most of these N-termini were represented exclusively by one or several dimethylated peptide(s) (74 % and 56 % in chromatophore and nucleus-encoded proteins, respectively), with the remaining N-termini represented by acetylated or both, dimethylated and acetylated peptide(s) (**Fig. 1B**).

### N-terminal Processing of Chromatophore-Encoded Proteins

For 87 of 132 (or 66 %) of the chromatophore-encoded proteins, canonical N-termini were found, i.e., the protein was not processed (26 proteins), only the iMet was removed (54 proteins) or both proteoforms were identified (7 proteins). For 27 of these proteins additional non-canonical N-terminal peptide(s) that did not match any known or predicted cleavage site were observed, indicating the presence of additional and presumably protease-generated proteoforms. 45 proteins were exclusively identified by non-canonical N-termini. 39 % of these non-canonical N-termini mapped within the first 20 aa and 62 % within the first 50 aa of the corresponding protein models (**Fig. 2A, Table S1**).

**Figure 2.**
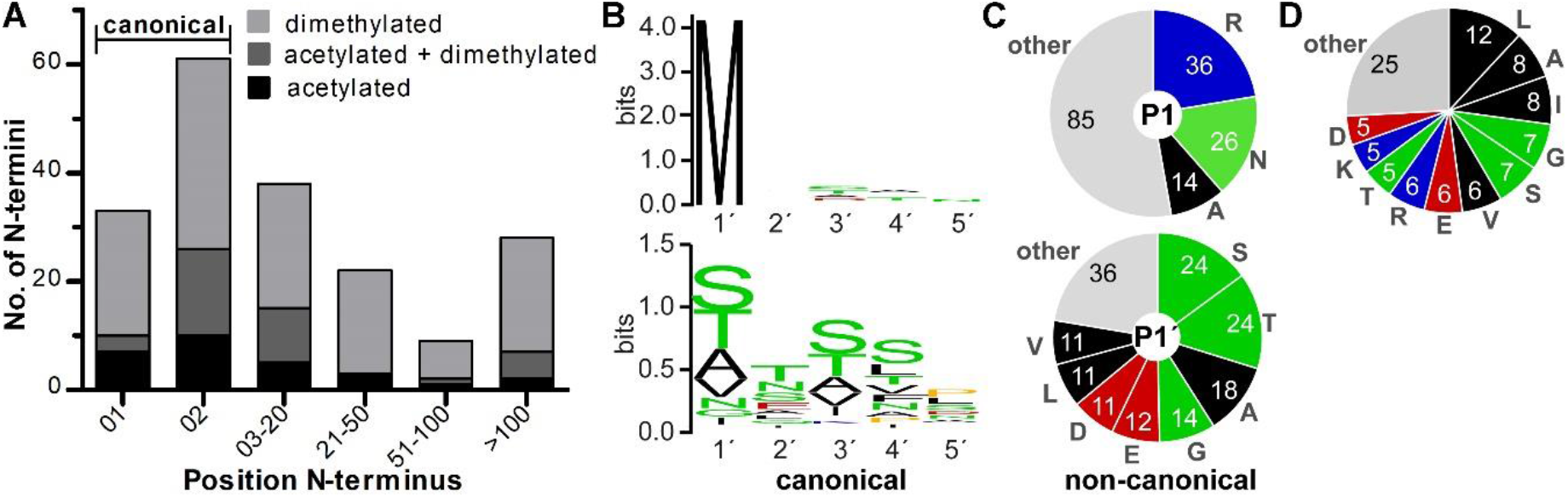
Position and amino acid composition of processing sites in chromatophore-encoded proteins. **(A)** Position of identified N-termini with respect to protein models. The number of proteins is depicted for which one or several N-terminal peptide(s) matching the respective protein model at the indicated positional ranges have been identified. Color code as in Fig. 1B. **(B)** Amino acid frequencies at canonical N-termini. The sequence logos are based on 33 unprocessed unique N-termini (top) or the 61 unique N-termini that result from iMet excision (bottom). Logos were created with weblogo (https://weblogo.berkeley.edu/logo.cgi). **(C)** Total numbers of the most frequent amino acids present at P1 and P1’
s positions of non-canonical N-termini. **(D)** Overall amino acid frequencies of the chromatophore-encoded proteome in percent, rounded to the full digit. **(B-D)** Black, hydrophobic; green, polar amino acids and glycine; red, negatively charged; blue, positively charged; yellow, proline.

Many of the proteins for which multiple N-termini were detected show particularly high abundance levels in the chromatophore as determined previously (Oberleitner et al., 2020); e.g., 10 N-termini have been identified for the phycobilisome linker polypeptide, 8 for the RubisCO large subunit (**Table S1**). However, there is no strict correlation between protein abundance and number of detected proteoforms as also other factors (protein lifetime, availability of trypsin cleavage sites, physicochemical properties of generated peptides, etc.) affect the number of detectable N-terminal peptides (Niedermaier and Huesgen, 2019). Often several N-termini identified for one protein were located within a range of only 10 aa, e.g., 6 of the 10 unique N-termini identified for phycobilisome linker polypeptide were located between position 19 to 27 (**Fig. S1**). This phenomenon termed “ragging” is frequently observed (Fortelny et al., 2015) and suggests either sloppy cleavage specificity of one protease, and/or additional processing by aminopeptidases (Rowland et al., 2015).

Some of these non-canonical peptides may certainly result from protein degradation *in vivo* or during sample preparation, other non-canonical termini likely represent biologically relevant proteoforms. This is supported by the occurrence of N-acetylation on non-canonical N-termini (**Fig. 2A**; **Table S1**). Approximately 38 % of the N-terminal peptides annotated as canonical or mapping to aa residues 3-20 of the corresponding protein models were acetylated, but only 20 % of the remaining N-terminal peptides (**Fig. 2A**). In some cases, the most abundant (often the canonical) and therefore likely most relevant N-terminus could be distinguished using peptide intensity as a proxy (see examples in **Fig. S1**). Notably, this is not an absolute measure of abundance, as peptide ionization efficiency depends on the peptide sequence and additional more abundant canonical N-termini may be present but not produce detectable tryptic peptides with a length between 8 to 40 aa (see **Table S1**). Furthermore, a few canonical N-termini might be misinterpreted as non-canonical due to mis-annotation of the correct translation initiation site (grey arrows in **Fig. S2B**). Finally, for four proteins of bin 21-100 the identified processing site matched a predicted signal peptide cleavage site (orange arrows in **Fig. S2B; Table S1**), among them are the thylakoid lumen proteins cytochrome c-550 (PCC0702) and the photosystem I reaction center subunit III (PsaF, PCC0760).

Generally, the distributions of amino acids occurring in position P1 (the residue preceding the identified N-terminus) and position P1’
s (the N-terminal residue of the identified N-terminus) are neither random nor do they reflect the overall frequencies of amino acids in the predicted chromatophore-encoded proteome (**Figs. 2B-D** and **S2**). The most common amino acids in P1 positions are methionine, arginine, asparagine, and alanine. Methionine in the P1 position is usually related to iMet excision in canonical N-termini (**Figs. 2B** and **S2**). In non-canonical cleavage sites, the most common amino acids in the P1 position are arginine (22 % of sites), asparagine, and alanine (together 25 % of sites; **Figs. 2C** and **S2**). In both, canonical and non-canonical N-termini, in the P1’ positions, serine, threonine, and alanine are the most common amino acids (**Fig. 2B**,**C**) and the respective N-terminal peptides are also highly abundant, pointing to a stabilizing effect of these N-terminal amino acids (**Fig. S2**).

### N-terminal Processing of crTP-Carrying Proteins and Common Features of the crTPs

Of the 76 nucleus-encoded proteins for which N-terminal peptides could be identified (**Fig. 1B, Table S1**), 37 proteins carry a crTP, 15 are short (<90 aa), 7 proteins are long but lack a crTP, and for the remaining 17 proteins only partial sequence information is available. Notably, 30 of the 37 crTP-carrying proteins were processed between alignment position 72 and 94, corresponding to aa position 37-69 in the protein sequences, which we therefore designate as processing region 1 (PR1) (**Fig. 3A**). As in chromatophore-encoded proteins, often multiple cleavage sites are found in close proximity, i.e., the generated proteoforms differ only by one to five amino acids (**Fig. 3A**). A correlation between the number of unique N-termini per protein and protein abundance as estimated by intensity of the corresponding N-terminal peptides (here), sum of peptide ion intensities in an independent MS analysis (Oberleitner et al., 2020) or transcript abundance (Nowack et al., 2016).

**Figure 3.**
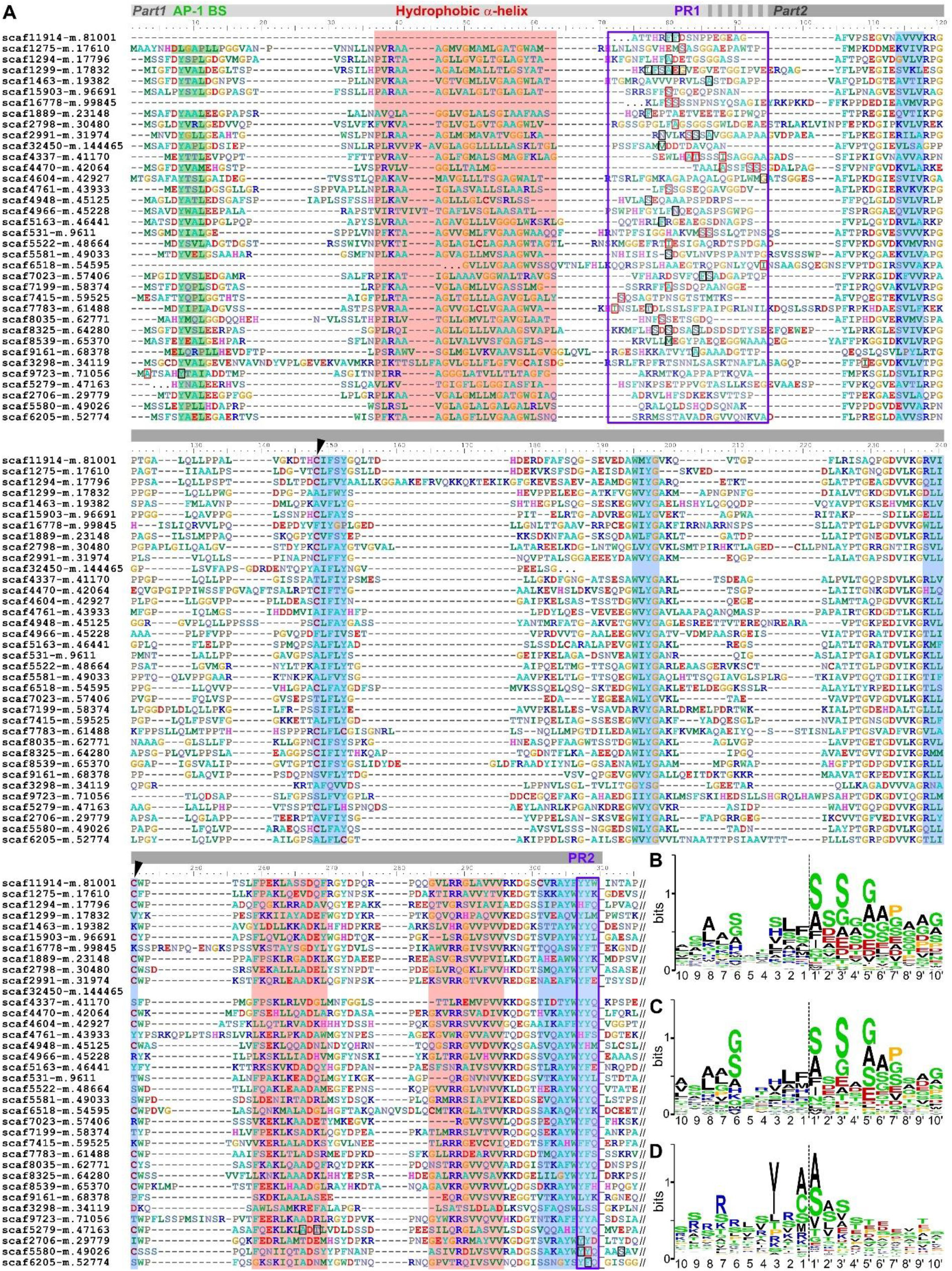
Position and amino acid composition of processing sites in crTP-carrying proteins. **(A)** Alignment of 36 crTPs for which N-termini were determined by HUNTER. Scaffold19070-m.107696 which contains a relatively divergent crTP sequence with long insertions was excluded from the alignment. Black and red rectangles surround the amino acids in the P1’ positions of identified dimethylated and acetylated (or both, acetylated and dimethylated) N-terminal peptides, respectively. The common processing regions 1 and 2 (PR1 and PR2) are framed in violet. Putative AP-1 binding sites (AP-1 BS) are shaded in green. Lacking sequence information is represented by dots and the C-terminal end of a crTP by an underscore. Areas containing conserved predicted α-helices or beta sheets are shaded in red and blue, respectively. The conserved cysteine pair is highlighted with black arrowheads. **(B-C)** Amino acid frequencies around cleavage sites (dashed lines) in crTP PR1. The sequence logo is based on **(B)** the 44 unique N-termini (with full length sequence information) from PR1 or **(C)** the 28 most N-terminal N-termini in PR1 when multiple N-termini per protein were found. **(D)** Amino acid frequencies around cleavage sites of the cTP of *Arabidopsis thaliana*. The sequence logo is based on the best-ranked N-termini of 162 nucleus-encoded stromal proteins according to Rowland et al., 2015. Logos were created and color coded as in Fig. 2B.

Although PR1 is located between two conserved regions in a poorly alignable region with no sequence conservation between individual crTPs, a preference for certain amino acids around the processing site becomes apparent (**Fig. 3B**). This pattern changes only little when considering only the most N-terminal peptide derived from PR1 of each protein (compare **Figs. 3B and 3C**). The region is overall rich in serine and glycine. Upstream of the processing site (P10 to P1) positive charges are more prevalent and phenylalanine is present at the P1 position in 25 % of all cases. Downstream of the processing site (P1’ to P10’) negative charges are frequent and this region is also comparably rich in serine, glycine, alanine, and proline. In the P1’
s position, serine is present in 41 % of all N-termini identified, followed by alanine (18 %), phenylalanine (12 %), and isoleucine (8 %). Notably, isoleucine is always acetylated, while serine and alanine are sometimes acetylated (30 % and 11 %, respectively), and phenylalanine is never acetylated. When multiple processing sites were identified in close proximity for a protein, the corresponding peptides usually differed in their relative intensity (**Table S1**). However, no obvious prevalence for a certain N-terminal amino acid, acetylation status or relative site position was observed.

When compared to cTP processing sites of proteins imported into plastids in *Arabidopsis* (**Fig. 3D**; Rowland et al., 2015), the glaucophyte alga *Cyanophora paradoxa* (Köhler et al., 2015), and the diatom *Thalassiosira pseudonana* that harbors complex plastids (Huesgen et al., 2013), some similarities become apparent. This includes for example the distribution of charges upstream and downstream of the processing site, the prevalence of serine and alanine at the P1’
s position, and an overall high frequency of serine (**Figs. 3B and D**). However, in contrast to the cTP processing site, glycine is overall more common around the crTP processing site and the occurrence of phenylalanine at the P1 and P1’ and isoleucine at the P1’ positions clearly distinguish the crTP from the cTP cleavage sites.

Interestingly, only for one crTP-carrying protein that is cleaved in PR1 (scaffold1294-m.17796; annotated as ‘kelch domain-containing protein’), additional N-termini downstream of the crTP were detected. These N-termini are located 60 and 62 aa downstream of the C-terminal end of the crTP in a region that shows no homology to other known proteins (**Table S1**). For six other crTP-carrying proteins no peptides derived from cleavage in PR1 but from other positions within the crTP were obtained, notably from regions of high sequence conservation (**Fig. 3A**). One crTP (in scaffold3298-m.34119, annotated as ‘Short-chain dehydrogenase Tic32’) is cleaved just a few aa downstream of PR1. Notably, this crTP lacks negatively charged amino acids in PR1 which typically can be found in P1’-P10’ positions. In one crTP (in scaffold9723-m.71056, annotated as ‘HIT-like protein’) solely the iMet is removed resulting in the only canonical N-terminus identified for a crTP-carrying protein and a second N-terminal peptide 5 aa downstream was identified but no additional N-terminal peptides from regions further downstream; this canonical N-terminus may be derived from an intact precursor protein before import. Only three proteins are processed at the C-terminal end of the crTP (alignment position 307-308, denoted PR2 in **Fig. 3A**), i.e., rather the expected position for full removal of a targeting sequence. One is annotated as ‘metal-dependent protein hydrolase’, the other two as ‘SDR family NAD(P)-dependent oxidoreductases’. In an attempt to identify more N-terminal peptides corresponding to cleavage in PR2, the *P. chromatophora* transcriptome database was queried a second time using a relaxed false discovery rate (FDR =0.05). Although this led to the identification of approx. 27 % more nucleus-encoded peptides and further three crTP-carrying proteins processed at PR1, no further PR2 N-termini were found.

Thus, surprisingly, for 33 crTP-carrying proteins no N-termini at the start of the functional protein were identified, suggesting that after cleavage in (predominantly) PR1 the C-terminal part 2 of the crTP remains attached to the N-terminus of the imported protein. Importantly, this finding is supported by mapping peptides identified in previous shotgun proteome analyses on the crTP alignment (**Fig. S3**). This analysis demonstrates that peptides identified in chromatophores derive not exclusively from the functional protein but at similar sequence coverage and peptide intensity levels from crTP part 2 (**Fig. S3**).

To find hints pointing towards a crTP-mediated import mechanism, we analyzed the crTP sequences with various bioinformatic approaches. Interestingly, the Eukaryotic Linear Motif resource (http://elm.eu.org/) identified the sorting signal YxxΦ (where “Y” stands for tyrosine, “Φ” for a bulky, hydrophobic, and “x” for any aa) at the N-terminus of all but two crTPs for which full-length sequence information is available (**Fig. 3A**). This motive is involved in protein sorting at the *trans* Golgi network (TGN) and is found in the cytoplasmic tail of membrane-spanning cargo proteins. The motive is recognized by the adaptor protein-1 complex (AP-1), a clathrin adaptor that couples cargo recruitment to coated vesicle budding (Park and Guo, 2014). Noteworthy, the two crTPs lacking the YxxΦ signal possess the motif [DE]LxxPLL instead which is not identical but very similar to the motif [DE]xxxL[LI] which represents an alternative sorting signal bound by AP-1 (Park and Guo, 2014) and to further motif variants (e.g., EAAAAPLL) that have been experimentally shown to bind AP-1 in human cells (Kozik et al., 2010). These putative AP-1-binding sites are followed by a conserved hydrophobic α-helix that is predicted in some but not all crTPs as a transmembrane helix (TMH) by the TMHMM algorithm (Krogh et al., 2001). PR1 lies downstream of the hydrophobic helix in a region that is mostly predicted as unstructured by IUPred2A (Mészáros et al., 2018). The rest of the crTP sequence contains several conserved secondary structural elements predicted by PROMALS3D (PROfile Multiple Alignment with predicted Local Structures and 3D constraints; http://prodata.swmed.edu/promals3d) (Pei et al., 2008) as well as a conserved cysteine pair that is present in half of the proteins analyzed (**Fig. 3A**) suggesting that part 2 of the crTP folds into a common 3D structure.

### N-terminal Processing of Short Imported Proteins

Of the 15 short imported proteins for which N-terminal peptides were identified here, six were detected by MS as imported proteins before, while the remaining nine represent new identifications (**Table S1**). Most of the 15 proteins lack a functional annotation but several match to ‘group 1 to 4’ short imported proteins described before (Oberleitner et al., 2020) (**Table S1**). Only scaffold27615-m.132528 is annotated as ‘carboxysome peptide A’. In line with the notion that these proteins lack obvious targeting signals at their N-terminus, for all short imported proteins only one N-terminal peptide has been identified usually corresponding to the canonical N-terminus. In most cases (10 out of 15 proteins) the iMet is removed, two proteins remain entirely unprocessed (including the carboxysome peptide A), and in one protein the first two aa are cleaved off (**Fig. 4A**). Interestingly, the only two short imported proteins that are processed by cleavage of more than two N-terminal aa, also represent the only proteins that possess a predicted TMH [i.e., ‘group 1 short imported proteins’ (Oberleitner et al., 2020)]. In both cases, cleavage occurs among the ten N-terminal aa between alanine and a negatively charged residue (**Fig. 4A**).

**Figure 4.**
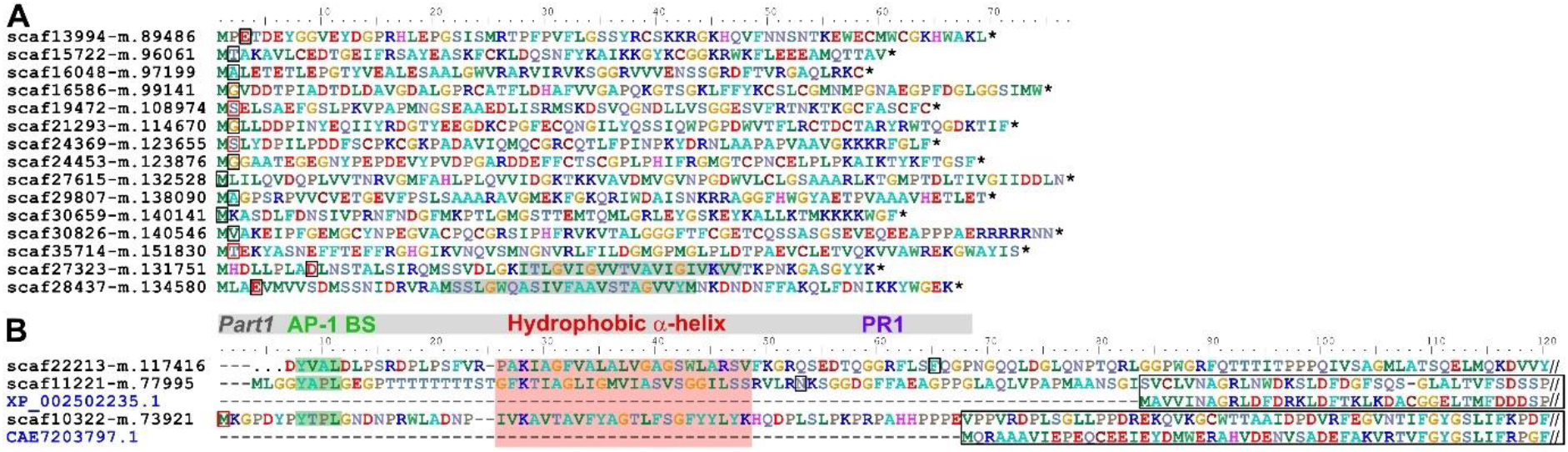
Position of processing sites in short (A) and unusual long (B) imported proteins. **(A)** Short imported proteins for which N-terminal peptides were identified. P1’ positions are represented as in Fig. 3A. Predicted TMHs in ‘group 1 short imported proteins’ are shaded in grey. **(B)** Three long imported proteins carry only crTP part 1 at their N-terminus (representation of sequence features as in Fig. 3A). If available, best blastp hits (NCBI) are shown in blue and their aa sequences aligned with the imported chromatophore protein (black frame). XP_002502235.1, predicted protein [*Micromonas commoda*]; CAE7203797.1, chac2 [*Symbiodinium microadriaticum*].

### N-terminal Processing of Other Nucleus-Encoded Proteins

Although the large majority of long nucleus-encoded proteins identified in the chromatophore samples carry a crTP, a small number of proteins >250 aa without a crTP was found before in chromatophore samples (Singer et al., 2017; Oberleitner et al., 2020). It is currently unclear whether these proteins represent contamination, are imported or are associated with the chromatophore surface. However, note that the outer chromatophore membrane is largely lost during chromatophore isolation (Oberleitner et al., 2020). Also in the current study, N-terminal peptides for seven long nucleus-encoded proteins that do not possess a crTP, have been identified (**Table S1**). Interestingly, inspection of the amino acid sequence of these proteins revealed for two of them (plus a third for which full-length sequence information is missing) an N-terminus resembling part 1 of the crTP (i.e. YxxΦ signal, hydrophobic α-helix, followed by an unalignable region; compare **Figs. 3A** and **4B**), whereas part 2 of the crTP is missing. Instead, in the two proteins for which homologous sequences are found in other organisms, part 1 is immediately followed by the conserved protein (**Fig. 4B**). For two of these proteins N-terminal peptides resulting from cleavage 4-16 aa downstream of the hydrophobic helix were obtained, while for the remaining one only a canonical N-terminus was identified (**Fig. 4B**).

## Discussion

### Characteristics of the Chromatophore-Encoded N-Terminome

N-terminal methionine excision is an evolutionary conserved co-translational process that occurs in eukaryotes as well as prokaryotes and organelles and is generally restricted to proteins that possess small, uncharged amino acids in their penultimate positions (Varland et al., 2015). However, the precise aa preferences vary between organisms (Bonissone et al., 2013). Here we found that 65 % of the canonical proteoforms synthesized in the chromatophore are targets of iMet excision, preferentially when serine, threonine, alanine, or valine are the penultimate residues (**Figs. 2B** and **S2A**). Responsible for iMet excision is likely the chromatophore-encoded protein PCC0019 that shows 62 % similarity to the Met-aminopeptidase MatC from *Synechocystis* sp. 6803 (sll0555; Atanassova et al., 2003; Drath, Baier, and Forchhammer 2009). Overall, the frequency and specificity of iMet excision observed in chromatophores is comparable to the ones in free-living cyanobacteria and plastids (Sazuka et al., 1999; Giglione et al., 2004; 2015; Bonissone et al., 2013; Rowland et al., 2015).

Functionally, iMet excision can link N-terminus identity to its *in vivo* stability by what is known as the “N-end rule pathway” (Dissmeyer et al., 2018). This mechanism regulating protein stability has emerged as an important regulator of developmental processes and responses to environmental cues in plants and existence of a prokaryote-type pathway in plastids has been proposed (Rowland et al., 2015; Dissmeyer et al., 2018). Further experimentation will be needed to determine to what extent also chromatophore proteostasis is governed by an N-end rule.

Furthermore, the data presented here revealed that almost half of the canonical N-termini of chromatophore-encoded proteins are acetylated (**Fig. 2A**), especially when exposing methionine, threonine, alanine, or valine (**Fig. S2A**). Although in cyanobacteria, N-terminal threonine, valine, serine, and alanine can be acetylated following iMet excision (Bonissone et al., 2013), the overall frequency of N-terminal acetylation is low in bacteria (<5 %; Soppa 2010; Kouyianou et al., 2012; Yang et al., 2014; Schmidt et al., 2016), but much higher for plastid-encoded proteins (Giglione and Meinnel, 2001; Huesgen et al., 2013; Rowland et al., 2015). Hence, the frequency of N-acetylation in the chromatophore is more comparable to plastids, suggesting the involvement of host-derived factors in this process. In line with this assumption, no chromatophore-encoded N-acetyltransferase could be identified. Nevertheless, also orthologs of the common eukaryotic ribosomal N-acetyltransferases (Linster and Wirtz, 2018) do not appear to be imported into chromatophores, i.e., they were identified only in whole cell lysates in MS-based proteome analyses (NatA, NatC; Singer et al., 2017; Oberleitner et al., 2020) and do not possess a crTP (NatA, NatB, NatC, NatD, NatE). An ortholog of the nucleus-encoded N-acetyltransferase NatG that has become specialized for post-translational N-terminal acetylation of nucleus but also plastid-encoded proteins in *Arabidopsis* (Dinh et al., 2015), is missing in *P. chromatophora*.

The functional consequences of N-terminal acetylation are context-dependent and include protein stabilization or conditional destabilization (Dissmeyer et al., 2018), protein folding or aggregation, subcellular localization, and enhanced protein-protein interactions (Varland et al., 2015; Linster and Wirtz, 2018). Finally, N-terminal acetylations can be involved in adjusting organelle function to the physiological state of the cell, as was shown for plant ATP-synthase subunit Ɛ (Hoshiyasu et al., 2013). In chromatophores, acetylation seems to have a stabilizing effect on proteins possessing N-terminal valine or isoleucine and a destabilizing effect on proteins showing methionine, serine, glycine or alanine at their N-termini, as judged from intensities of the corresponding N-terminal peptides (**Fig. S2**).

Besides the canonical N-termini, with 63 % of the N-termini detected for chromatophore-encoded proteins, a relatively large number of non-canonical N-termini were found in this study. High rates of non-canonical proteoforms have also been reported in studies using similar methods (i.e., TAILS or COFRADIC) for determination of the N-terminome of plastids (38 %, Huesgen et al., 2013; 40 %, Rowland et al., 2015) or photosynthetic bacteria (35-60 % depending on cell fraction, Kouyianou et al., 2012). As observed in chromatophores (**Fig. 2**), many of these N-termini are generated by cleavage C-terminal to arginine (or asparagine) residues and create N-termini starting with threonine or serine (Kouyianou et al., 2012; Rowland et al., 2015; Berry et al., 2017). These non-canonical N-termini are commonly classified as unknown proteolytic cleavage products or degraded proteins and excluded from further analyses.

However, ∼20 % of the non-canonical N-termini identified here are acetylated and at least a few seem to represent processed clients of the Sec pathway [likely cleaved by the chromatophore-encoded signal peptidase (PCC0690) that shows 42 % similarity to LepB1 from *Synechocystis* sp. 6803 (sll0716)]. Thus, a number of non-canonical N-termini detected for chromatophore-encoded proteins clearly represent N-termini of functionally relevant proteoforms. Further known mechanisms for the generation of functional proteoforms from one pre-protein are for example N-terminal trimming of several amino acids by exo- or endo-peptidases, zymogen activation via cleavage of a pro-peptide, etc. (Lange and Overall, 2013; Perrar et al., 2019). Thus, the catalogue of proteoforms generated here for chromatophore-encoded proteins represents a valuable informational resource for detailed functional studies of specific proteins in the future.

### N-Terminal Processing of the crTP and Proposed Model for Import of crTP-Carrying Proteins into the Chromatophore

The identification of putative AP-1 binding sites at the N-terminus of all crTPs followed by a predicted TMH combined with the earlier detection of the short imported protein PsaE in the Golgi (Nowack and Grossman, 2012) prompted us to propose the following model for crTP-mediated protein trafficking via the Golgi (**Fig. 5**). In this model, the N-terminal TMH of the crTP part 1 triggers co-translational import of the protein into the ER via the signal recognition particle (SRP) system (**Fig. 5B**). The TMH is released by sideward opening of the Sec61 channel and anchors the protein in the ER membrane in the N-terminus out, C-terminus in conformation while travelling to the *trans* Golgi network (TGN). There, the sorting signals YxxΦ and [DE]LxxPLL are recognized by the clathrin adaptor protein complex AP-1, which recruits clathrin units (and other factors) to the membrane initiating vesicle formation (**Fig. 5C**). Along with the crTP-carrying cargo proteins, also dedicated v-SNAREs are incorporated into the vesicle membrane. Once the vesicle pinched off the Golgi, the clathrin coat is lost. As soon as a vesicle approaches the chromatophore, v-SNAREs bind to their t-SNARE counterparts in the chromatophore outer membrane. Vesicle fusion is initiated, and crTP-carrying cargo proteins enter the chromatophore intermembrane space (IMS) (**Fig. 5D**). Here, either, a specific endopeptidase cleaves part 1 of the crTPs releasing the soluble protein into the IMS and exposing part 2 of the crTP (scenario 1) or the hydrophobic helix is pulled out of the outer membrane, maybe stabilized by chaperones and mediates also translocation across the chromatophore inner membrane (scenario 2). Thus, in scenario 2, part 1 of the crTP would be cleaved only following translocation across the inner membrane.

**Figure 5.**
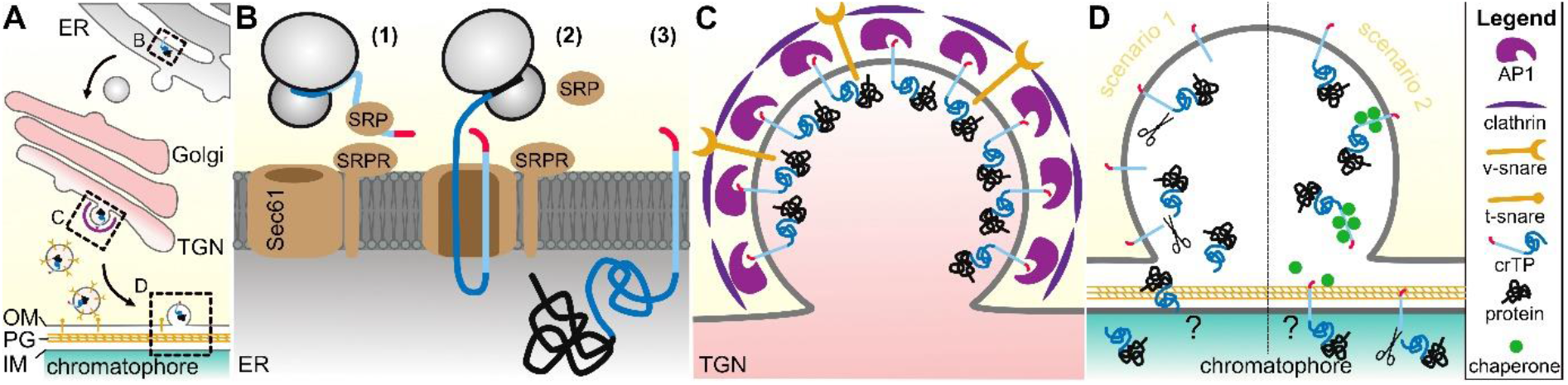
Hypothetic model for crTP-mediated protein import into the chromatophore. **(A)** Overview; **(B-D)** details as indicated in A. Red, putative AP-1 complex binding motifs (YxxΦ and [DE]LxxPLL); light blue, TMH of crTP part1; dark blue, part 2 of the crTP; black, functional protein. OM, outer chromatophore membrane; PG, peptidoglycan; IM, inner chromatophore membrane. For details see the text.

We observed a remarkable similarity between aa composition and charge distribution around PR1 localized crTP cleavage sites and the cTP cleavage site in plants and algae. In plants, cleavage of cTPs on thousands of different proteins is achieved by a single nucleus-encoded M16-type metallopeptidase (Stromal Processing Peptidase, SPP) that recognizes rather charge distribution and structural features around the cleavage site than aa sequence (Richter and Lamppa, 1998). In *P. chromatophora*, a putative M16-type processing peptidase with similarity to SPP or related cyanobacterial proteins (Richter et al., 2005) could not be identified among chromatophore-encoded or imported proteins. Thus, identity of the crTP processing peptidase as well as its subcellular localization remain unknown.

Remarkably, >80 % of the crTP-carrying proteins for which N-terminal peptides were identified appeared to retain part 2 of the crTP at the N-terminus of the imported protein (**Fig. 3A**). In chloroplasts, a free N-terminus generated by cleavage of the complete targeting peptide is required for the correct function and subcellular localization (e.g., the thylakoid lumen) of many proteins (Richter et al., 2005). Nevertheless, survival of crTP part 2 seems to be supported also by detection of abundant part 2-derived MS spectra in independent MS analyses (**Fig. S3;** Singer et al., 2017; Oberleitner et al., 2020). An alternative interpretation could be that part 2 crTPs survive as independent proteins after cleavage from the mature imported protein, whereas most tryptic peptides derived from the true N-terminus of the mature protein are disguised by a common characteristic rendering them invisible to detection by LC-MS/MS (e.g., posttranslational modifications, length of the peptides outside of the detection range, bad ionization properties, etc.). However, close inspection of the transition region between crTP and conserved domains of the functional proteins did not yield any obvious reason for a possible inability to detect corresponding tryptic peptides. And also in our previous shotgun proteome analyses, peptides that span this transition zone were detected (**Fig. S3C**), supporting the disposition of crTP part 2 at the N-terminus of the mature protein rather than its survival as independent protein.

Only for three proteins complete removal of the crTP was found (i.e., cleavage in PR2 in **Fig. 3A**). It is possible that only those proteins that require a free N-terminus for their correct function or subcellular localization (e.g., the thylakoid lumen) acquired a PR2 cleavage site. However, neither protein annotation nor aa sequence around PR2 cleavage sites in processed vs. non-processed crTPs (such as Sec or Tat secretion signals following the conserved crTP sequence) clearly support this hypothesis. Notably, our previous shotgun proteome analyses identified crTP part 2-derived peptides also for proteins for which HUNTER detected cleavage sites outside of PR1 (**Fig. S3**). This finding suggests that N-termini outside of PR1, which are mainly represented by dimethylated peptides, might represent degradation products arising from regular protein turnover.

There are still many further questions associated with the import of crTP-carrying proteins. Maybe the most central one is how the crTP (or part 2 of the crTP) mediates protein translocation across the inner chromatophore membrane. When released into the chromatophore IMS following passage of the Golgi, crTP part 2 is likely fully folded, and potentially stabilized by a disulfide bond between the paired cysteine residues in half of the proteins (arrowheads in **Fig. 3A**). Whether these proteins are unfolded by unfoldase activity and threaded through an import channel of unknown identity in the inner membrane or transported in the folded state by an unknown mechanism is currently unclear. In any case, a striking observation is that both, the canonical N-terminus of the crTP as well as the processed N-termini resulting from crTP cleavage in PR1 are dominated by negatively charged amino acids (average charges of -1.7 or -2.0 at pH 7.0 over the first 20 aa, respectively; calculated from all 32 and 49 available sequences identified here). This differs remarkably from proteins being post-translationally imported into mitochondria and plastids as well as proteins being transported across bacterial membranes by the Sec or the Tat system, which all typically feature positively charged pre-sequences (Garg and Gould, 2016).

Finally, an interesting observation in this study was that some long chromatophore-targeted proteins without a full length crTP seem to carry an isolated crTP part 1 at their N-terminus. Determination of the subcellular localization of these proteins might be key to distinguish between different possible import mechanisms as scenario 1 in **Fig. 5D** would predict these proteins to localize to the chromatophore IMS while scenario 2 would rather predict these proteins to completely translocate across the chromatophore inner membrane.

## Conclusions

In sum, this work yielded a detailed picture of N-terminal processing and modification of chromatophore-localized proteins in *P. chromatophora*. The level of N-terminal acetylation in chromatophores, possibly by nuclear factors, is more comparable to N-acetylation patterns in plastids than in cyanobacteria and might contribute to the adjustment of chromatophore performance to the physiological state of the cell.

The onset of protein import into a recently established organelle is at the heart of the transformation from bacterial endosymbiont to genetically integrated organelle. Thus, a detailed understanding of protein import into the chromatophores of *Paulinella* will provide precious mechanistic insights into the process of organellogenesis that cannot be gained from evolutionary more derived systems. Our study revealed the fate of the crTP upon import into the chromatophore and enabled the proposal of a model for crTP-mediated protein import. The proposed mechanism differs fundamentally from protein import mechanisms known from ancient eukaryotic organelles and will be helpful to guide future experimentation using biochemical approaches, including *in vitro* import assays.

## Material and Methods

### Cultivation of *P. chromatophora* and Isolation of Chromatophores

*P. chromatophora* CCAC0185 (axenic culture) was grown and chromatophores isolated essentially as described before (Singer et al., 2017). In brief, cells were washed three times with isolation buffer [50 mM HEPES pH7.5, 250 mM sucrose, 125 mM NaCl, 2 mM EGTA, 2 mM MgCl_2_, protease inhibitor cocktail (Roche cOmplete)]. Cells were broken in a cell disruptor (Constant Systems) at 0.5 kbar and intact chromatophores isolated on a discontinuous 20-80 % Percoll gradient. To increase yield, the pellet at the bottom of the gradient was subjected to another round of cell disruption and chromatophore isolation. To increase purity, pooled chromatophores were re-isolated from another, fresh Percoll gradient. All steps were carried out at 4°C.

### Preparation of Chromatophore Lysate

Chromatophore lysate was prepared in triplicates from approximately 6 × 10^6^ chromatophores each. Immediately following isolation, chromatophores were washed in 500 µl wash buffer (100 mM HEPES pH 7.5, 5 mM EDTA) and then resuspended in 300 µl lysis buffer (100 mM HEPES pH7.5, 5 mM EDTA, 6 M Guanidine-HCl). ∼200 µl acid-washed glass beads (0.4-0.6 mm diameter) were added and chromatophores were lysed by vortexing for 5 min at room temperature. Then, the mixture was heated for 10 min at 95°C, vortexed again for 5 min, and transferred to a new tube without the glass beads. Glass beads were washed with 100 µl lysis buffer and fractions were combined. Insoluble material and residual glass beads were removed by centrifugation at 20,000 x g for 10 min, the clear lysate was transferred to a new tube, and the protein concentration determined by 660 nm Protein Assay (Pierce). Lysates, each containing 200-250 µg protein, were frozen in liquid nitrogen and stored at -80°C until further processing. Protein LoBind tubes (Eppendorf) and epT.I.P.S. LoRetention tips (Eppendorf) were used during all steps. Protease inhibitor cocktail (Roche cOmplete) was added to all buffers used.

### HUNTER

Protein disulfide bonds in chromatophore lysates were reduced by the addition of DTT to a final concentration of 10 mM and incubation at 37°C for 30 min. Reduced cysteine residues were carbamidomethylated by the addition of chloroacetamide to a final concentration of 50 mM and incubation at RT for 30 min in the dark to prevent reformation of disulfide bonds. Chloroacetamide was quenched by further addition of DTT to 50 mM final concentration and incubation at RT for 20 min. Reduced proteins were now purified from the lysate samples using SP3 magnetic beads (1:1 mixture of SpeedBead Magnetic Carboxylate Modified Particles 65152105050250 and 45152105050250, GE Healthcare). To enable protein binding to the beads, ethanol was added to a final concentration of 80 % (v/v). Then, 1 µl bead suspension was added per 20 µg of protein and the suspension was shortly mixed in a sonication bath and incubated on a rotary shaker at RT for 20 min. Next, liquid was removed in a magnetic rack, the beads washed twice with 400 µl of 90 % (v/v) acetonitrile under short sonication, and finally, beads were air dried. Proteins were detached from the beads by the addition of 30 µl 100 mM HEPES buffer (pH 7.4) but the beads remained in the samples during the next steps.

Both N-terminal alpha-amines and lysine side chain amines were now labeled via reductive dimethylation by the addition of formaldehyde isotope CD_2_O to a final concentration of 30 mM and cyanoborohydride (NaBH_3_CN) to 15 mM. The samples were incubated at 37°C for 1 h. Another 30 mM formaldehyde and 15 mM cyanoborohydride were added and incubated as before to ensure complete labeling. Subsequently, the reaction was quenched in 500 mM Tris buffer (pH 7.4) at 37°C for 30 min. Proteins were again bound to the SP3 magnetic beads still present in the samples and purified as described earlier.

Proteins were now subjected to tryptic digest to generate peptides that can be identified via mass spectrometry. 30 µl of digest buffer (100 mM HEPES pH7.4, 5 mM CaCl_2_) and 1 µg of MS approved porcine pancreas trypsin (Serva) per 100 µg of protein were added and the mixture was incubated at 37°C and 1,200 rpm overnight inside a thermomixer with heated lid. Notably, cleavages can only occur C-terminal to arginine as cleavages at lysine residues are blocked due to dimethylation of lysine side chains.

Free alpha-amines on trypsin generated internal and C-terminal peptides were now hydrophobically tagged using undecanal. Ethanol and undecanal were added to final concentrations of 40 % (v/v) and 50 µg per 1 µg protein, respectively, and mixed gently by inverting. The tagging reaction was started by the addition of cyanoborohydride to 30 mM and the samples were incubated at 37°C and 1,200 rpm for 1 h. HR-X spin columns (Macherey-Nagel) containing hydrophobic polystyrene-divinylbenzene copolymer were used to deplete undecanal-labeled peptides from the samples. Sample volume was filled up to 400 µl with 40 % (v/v) ethanol and magnetic beads were washed three times. The samples were loaded on HR-X columns (activated beforehand by two additions of 400 µl methanol and equilibrated 2 x with 400 µl 40 % (v/v) ethanol), centrifuged for 1 min at 50 x g, and the eluate was collected. Another 400 µl 40 % (v/v) ethanol were loaded on the columns to elute remaining peptides by centrifugation as described before. Both fractions were combined and the solvent evaporated in a vacuum centrifuge at 60°C to complete dryness.

Enriched N-terminal peptides were resuspended in 40 µl of 0.1 % (v/v) formic acid (1 < pH < 3) and loaded on self-packed double layer C18 stage tips (activated with 20 µl of a mixture of 50 % acetonitrile and 0.1 % formic acid and equilibrated with 40 µl 0.1 % formic acid). Bound peptides were washed with 50 µl 0.1 % formic acid and eluted with 20 µl of a mixture of 50 % acetonitrile and 0.1 % formic acid. The solvent was evaporated in a vacuum centrifuge at 60°C. Peptides were resuspended in 15 µl 0.1 % formic acid and the peptide concentration was determined photometrically with a NanoDrop 2000c spectral photometer (Thermo Fischer Scientific) at 280 nm against a series of peptide standards.

### Mass Spectrometric Analysis and Protein Identification

Samples were analyzed on a nano-HPLC (Ultimate 3000 nano-RSLC) system equipped with a reverse-phase trap column (2 cm µPAC trapping column, PharmaFluidics) and a reverse-phase analytical column (50 cm µPAC column, PharmaFluidics). Peptides were eluted with a gradient from 2 to 30 % of solution B for 90 min (A: H2O + 0.1 % formaldehyde, B: acetonitrile + 0.1 % formaldehyde) and online introduced into a high-resolution Q-TOF mass spectrometer (Impact II, Bruker) using a nano-spray ion source (CaptiveSpray, Bruker) as described (Beck et al. 2015). Data was acquired in line-mode in a mass range from 100 to 1400 m/z at an acquisition rate of 10 Hz using the Bruker HyStar Software (v5.1, Bruker Daltonics). The top 14 most intense ions were selected for fragmentation. Fragmentation spectra were dynamically acquired with a target TIC (= total ion current) of 25k and a minimal frequency of 5 Hz and a maximal frequency of 20 Hz. Fragment spectra were acquired with stepped parameters, each with half of the acquisition time dedicated for each precursor: 80 µs transfer time, 7.5 eV collision energy, and a collision radio frequency (RF) of 1500 Vpp or 120 µs transfer time, 10 eV collision energy, and a collision RF of 1700 Vpp.

Obtained MS data was queried in a database search using MaxQuant v1.6.8.0 (Tyanova et al., 2016) with the standard settings for Bruker Q-TOF instruments. Searches were carried out using 60,108 sequences translated from a *P. chromatophora* transcriptome and 867 sequences derived from translated chromatophore ORFs (Singer et al., 2017). A database containing common contaminants, embedded in MaxQuant, was also included in the query. A decoy database was appended with the “revert” option. For HUNTER queries dimethylation of lysines and protein N-termini was set as fixed label (+32.0564 Da due to CD2O). Digestion mode was changed to semispecific (free N-terminus) ArgC and Oxidation (M), acetylation (peptide N-term) and Glu/Gln -> pyro-Glu were set as variable modifications, while carbamidomethylation of cysteines was set as fixed modification. Requantify option was enabled and maximal peptide length for unspecific searches was set to 40 aa. The minimal number of ratio count was set to 1. N-terminal peptides identified by MaxQuant searches were further validated and annotated using MANTI (Demir et al., 2021).

### Accession numbers

All MS-based proteomics data have been deposited to the ProteomeXchange Consortium via the PRIDE (Perez-Riverol et al., 2019) partner repository with the dataset identifier PXD028527 (reviewer login; password: kcoinut8).

## Acknowledgments

This study was funded by the Deutsche Forschungsgemeinschaft CRC 1208 project B09 (to E.C.M.N.).

